# Light-dependent conversion of body color change pattern during zebrafish development

**DOI:** 10.64898/2026.11.03.764145

**Authors:** Jiarun Zhang, Kazuhiko Takemae, Daisuke Kojima

**Author notes:** Correspondence: Daisuke KOJIMA.

## Abstract

Zebrafish (*Danio rerio*) shows body color change in response to light: when light comes, it whitens body color. Our previous study has shown that larval zebrafish as early as 5 days post-fertilization (dpf) have this capacity fully developed whereas 2-dpf larvae exhibit light-induced darkening in sharp contrast (Shiraki *et al*. 2010). Here, we discovered that the conversion in body color change pattern between 2 dpf and 5 dpf is light-dependent: larvae reared in darkness exhibited light-induced darkening of body color, reminiscent of 2-dpf larvae. We also found that a 14-h light exposure on 3 or 4 dpf is sufficient to trigger this conversion, and that the conversion by light on 4 dpf still occurred after ablation of most *exorh*-expressing pineal photoreceptor cells. The dark-reared larvae had significantly lower transcription levels of *pmch* and *pmchl*, which encode melanin-concentrating hormone precursors, than the LD-reared larvae. However, the levels in larvae that had undergone the conversion (3-L and 4-L) were not significantly different from those in dark-reared larvae, indicating that the reduced *pmch/pmchl* expression alone does not account for the conversion. In conclusion, light exposure in rearing is indispensable for the maturation of body color change ability in larval zebrafish through a regulatory mechanism independent of *exorh*-expressing pineal photoreceptors.

## Introduction

Zebrafish, as early as at larval stages, shows body color adaption to environment. When they are placed in the bright background, their body color becomes transparent. When they are in dark environment, correspondingly, they darken themselves. This ability is helpful for predator avoidance and categorized into physiological color changes in fish since it is done in a short period, involving dispersion or aggregation of melanin in melanophores.

The mechanism of body color change in teleost is being explored and it is believed that both nervous pathway and endocrine pathway are involved to transmit environmental hues to melanophores. On the one hand, sympathetic innervation plays a critical role in transmitting the signals from central nervous system to melanophores (Fujii, 2000). On the other hand, some hormones contribute to the dispersion or aggregation of melanin, such as melanin-concentrating hormones (MCH) that is secreted from the hypophysis and functions by aggregating melanin (Berman et al., 2009). Particularly, in zebrafish, *pmch* and *pmchl*, encoding MCH peptides, are expressed in lateral tuberal nucleus (NLT) (Berman et al., 2009) and MCH receptor *mchr2* is expressed on the skin of larval zebrafish (Madelaine et al., 2020). The dramatic increase of *pmch* and *pmchl* transcripts in white background (Berman et al., 2009) and *mchr2* mutants’ failure in whitening in bright environment (Madelaine et al., 2020), suggested that MCH induces aggregation of melanin in white background.

Some studies highlight that eye is essential for the background adaptation, but photoreceptors (rods or cones) are not. European minnows with the eyes removed failed to show whitening to light (Frisch, 1911) . *Lakritz* mutant zebrafish, in which most of retinal ganglion cells are not generated, failed to whiten in brightness, either (Neuhauss et al., 1999; Kay et al., 2001). Interestingly, in a forward genetic screening, 6 out of 7 groups of zebrafish mutant lines lacking most of photoreceptors showed a normal background adaptation (Muto et al., 2005), suggesting that other photosensitive cells in the retina might be involved or compensate for the loss of photoreceptors in background adaptation. Indeed, zebrafish has far more opsins than human beings and mice (Davies et al., 2015). By 5 days-post-fertilization (dpf), some retinal neurons other than cones and rods, such as retinal ganglion cells and bipolar cells, express multiple non-visual opsins (Davies et al., 2011; Matos-Cruz et al., 2011). Therefore, those neuron population in larval zebrafish may be candidate photosensitive cells that sense the brightness information for body color change.

The role of pineal in background adaptation is subtle. As mentioned above, removal of eyes abolishes the light-induced whitening (Frisch, 1911), which implied that pineal itself alone is not sufficient for this type of body color change. However, the *flh* mutant of zebrafish, in which the pineal neurogenesis is shut down but the eyes are remained morphologically intact (Masai et al., 1997), was reported to exhibit darker body color on white background and reduced expression of both *pmch* and *pmchl* (Zhang et al., 2010), suggesting that Agouti proteins secreted from pineal are essential for the whitening (Zhang et al., 2010). Thus, the pineal gland could be also highly important in light-induced whitening of body color in larval zebrafish, although it may not contribute to the light sensing reaction itself of body color change.

Our previous study (Shiraki et al., 2010) revealed a drastic conversion of light-induced body color change pattern during larval zebrafish development: Larval zebrafish on 2 dpf show a darkening of body color change in response to light, while on 5 dpf, they whiten the body color to light. This conversion may be explained by developmental maturation of neural and/or endocrine mechanism for light-induced whitening between these two stages, as 3-dpf and 4-dpf larvae exhibit a mixed pattern of light-induced darkening and whitening (Shiraki et al., 2010). Furthermore, 5-dpf larvae without eyeballs show a darkening of body to light (Shiraki et al., 2010), similar to the result show in minnows (Frisch, 1911), whereas isolated tails from 2-dpf larvae show a light-induced dispersion of melanin, i.e., darkening (Shiraki et al., 2010). Thus, the whitening is mediated by ocular photoreception and its downstream neural transmission while the darkening might be dependent on the local photoreception in dermal melanophores.

Dark rearing has been reported to have significant impacts on development of zebrafish and other vertebrates. In larval zebrafish, rearing in constant darkness results in loss of circadian rhythms of melatonin content (Kazimi and Cahill, 1999) as well as pineal gene expression of arylalkylamine-N-acetyltransferase (*aanat2*), a critical enzyme for melatonin synthesis (Ziv and Gothilf, 2006). Dark-reared zebrafish embryos and larvae also exhibit a delay in neurogenesis of habenular nuclei, which is attributed to reduction in melatonin accumulation (de Borsetti et al., 2011), reduced hatching rate, smaller body size and diminished feeding behavior (Villamizar et al., 2014). In mice, light activates the retinal wave in the second postnatal week and thereby facilitates retinotopic maps in the thalamus, while their refinement is reduced by dark rearing prior to eye opening (Tiriac et al., 2018). Dark rearing also impairs maturation of GABAergic circuits in the visual cortex of rats (Morales et al., 2002). These accumulated evidence for notable influences of light deprivation on vertebrate neural and endocrine systems prompted us to ask whether and how dark rearing affects the light-induced body color change ability in larval zebrafish.

Here, we report that light in rearing is indispensable for the developmental conversion of body color change pattern in larval zebrafish and that light exposure on 4 dpf is sufficient for inducing this conversion.

## Results

### Dark-reared 5-dpf larvae showed a light-induced darkening of body color

We have previously made a preliminary observation that dark-reared zebrafish larvae show darker body color under light illumination than those reared in the normal light-dark (LD) cycles. To figure out whether light deprivation in rearing has any effect on body color change ability, we reared larval zebrafish in the dark (DD) or under LD cycles until 5 dpf and then examined the body color change in response to light (Figure 1A). In the measurement on 5 dpf, the larvae were first dark adapted for 45 min, then exposed to light illumination for 30 min, and finally kept in the dark again for 45 min. In the LD-reared larvae (Figure 1B), the melanin index, which indicated the body color darkness, decreased upon light illumination (i.e., light-induced whitening) and then increased in the following dark period, being consistent with our previous report (Shiraki et al., 2010). In contrast, the DD-reared larvae exhibited an opposite pattern of body color change, i.e. light-induced darkening (Figure 1C), being reminiscent of body color change pattern of 2-dpf larvae (Shiraki et al., 2010). To extract metrics describing the difference in light-induced body color change between the two conditions, we calculated the area under curve (AUC), the primary metric defined in Materials and Methods, in the whole light phase (Figure 1D): The AUCs of DD-reared larvae showed a positive value in average and were significantly larger than those of the LD-reared (Figure 1E). We also compared the melanin index changes in light phase for 15 min or 30 min from light onset between the two conditions (Figure 1F, G): the DD-reared larvae showed 4% and 2% increase in melanin index during the 15 min and 30 min, respectively, which were significantly larger than the LD-reared (5% and 6% decrease, respectively). These results quantitatively demonstrated that light deprivation during zebrafish development converted the pattern of body color change from light-induced whitening to light-induced darkening.

**Figure 1.**
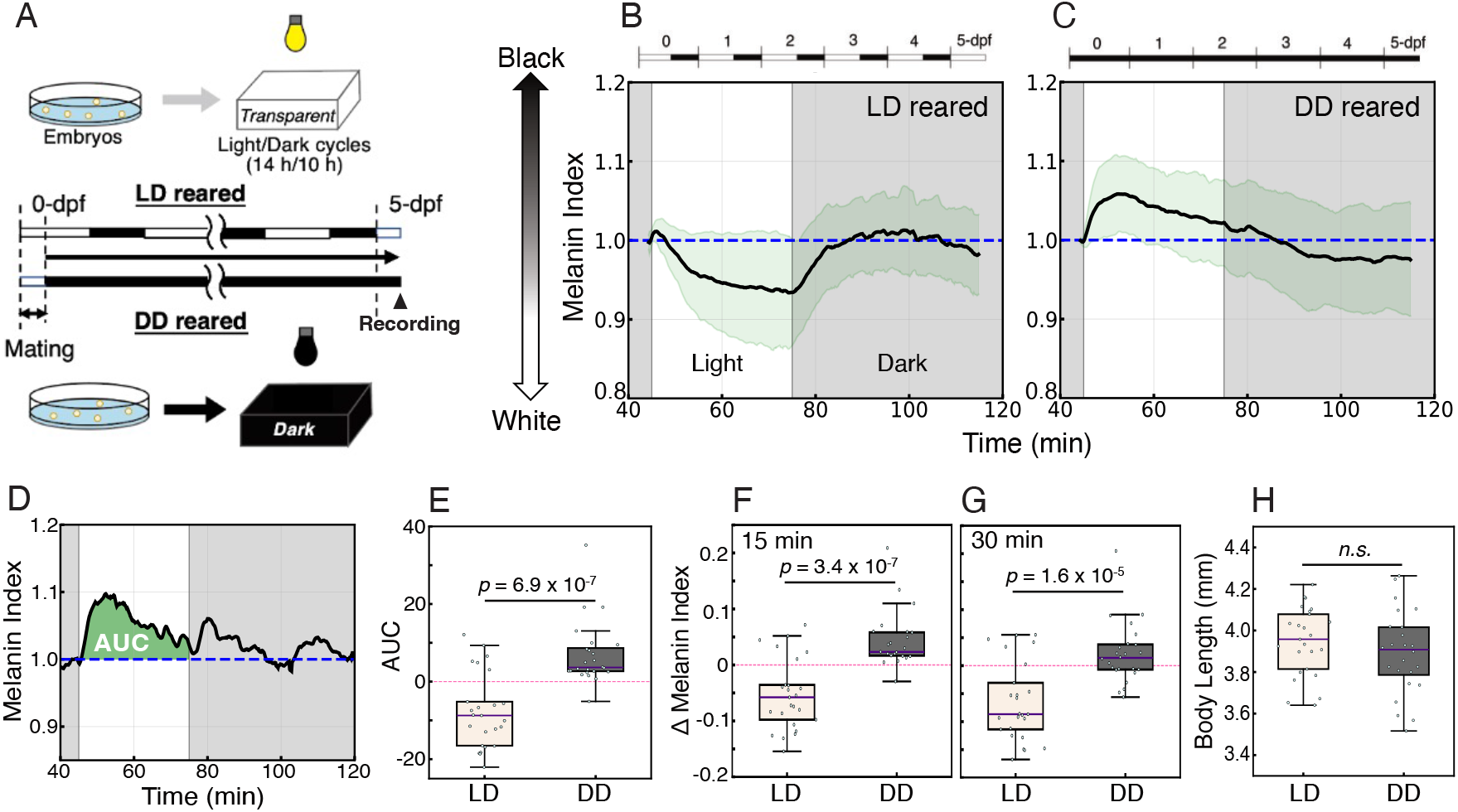
LD- and DD-reared larvae showed distinct patterns of light-induced body color change to light on 5 dpf. (A) Schematic drawing of LD- or DD rearing of larval zebrafish. (B, C) Time courses of melanin index in 5-dpf recordings of LD-reared individuals (B, n = 25) and DD-reared individuals (C, n = 24). Mean values of melanin index (± SD, grey shadow) are indicated. (D) An example of melanin index time course of DD-reared larva where the area under curve (AUC) is indicated in green. AUC is the sum of (MI − 1.0) over the 30-min light period; MI time courses are shown after smoothing (7-frame moving average) and normalization to MI = 1.0 at light onset (see Materials and Methods). (E) Comparison of AUC between the two conditions. (F, G) The melanin index changes of LD and DD group in 15 minutes (F) or 30 minutes (G) starting from light onset. (H) Comparison of body length between the two conditions. The *p*-values of Welch’s two sample *t*-tests between LD- and DD-reared group are indicated.

Note that, as shown in Figure 1C, light-induced darkening of 5-dpf dark-reared larvae was only detected in the first 10 min of light phase in contrast to the 2-dpf LD-reared larvae which exhibited darkening in the whole light phase (Shiraki et al., 2010). This observation suggested that light deprivation does not completely remove the ability of light-induced whitening but instead causes retardation or aberration in formation of neural circuit in charge of light-induced whitening. It should be also noted that dark-reared larval zebrafish exhibited a lower hatching rate, lower body size and higher fatality (Villamizar et al., 2013, 2014), which might reflect delay in development and be in relation to the light’s regulation to the cell cycle (Dekens et al., 2003). To exclude a possibility that those reported phenotypes might affect our results, we conducted a post-hoc analysis of the body size of the individuals, and revealed no significant difference between LD- and DD-reared larvae (Figure 1H).

### A period of light on 3 or 4 dpf gave ability of light-induced whitening on 5 dpf

Next, we aimed to figure out the timing when light is involved in the conversion in body color change because such information could help us to explore the mechanism of this light-dependent conversion. Firstly, we examined the timing of LD cycles in rearing required for the whitening of body color on 5 dpf. To this end, larval zebrafish were subjected to four conditions: LD (LD_5_DD_0_), LD_4_DD_1_, LD_3_DD_2_, and LD_2_DD_3_ (Figure 2A-D), where LD_n_DD_5-n_ meant that the light/dark cycles lasted for n times from the beginning followed by placing in the dark for 5-n days. On 5 dpf, LD- and LD_4_DD_1_-reared larvae presented light-induced whitening of body color (Figure 2A, B), but the other two groups, LD_3_DD_2_- and LD_2_DD_3_-reared, behaved differently with huge variations during the light phase (Figure 2C, D). On average, the latter two groups showed a transient darkening of body color within the first ∼5 minutes during the light phase and then turned to the baseline level of melanin index on average with a relatively large variation (Figure 2C, D), looking like intermediate between LD and DD groups (Figure 1B, C). These differences were also supported by quantitative comparisons of AUC and melanin index change during light exposure (Figure 2E-G), though the difference was not so significant in the melanin index changes in light phase for 15 min from light onset (Figure 2F). These results indicated that LD-cycle rearing lasting until 3 dpf (but not until 2 dpf) was sufficient to obtain ability of light-induced whitening on 5 dpf.

**Figure 2.**
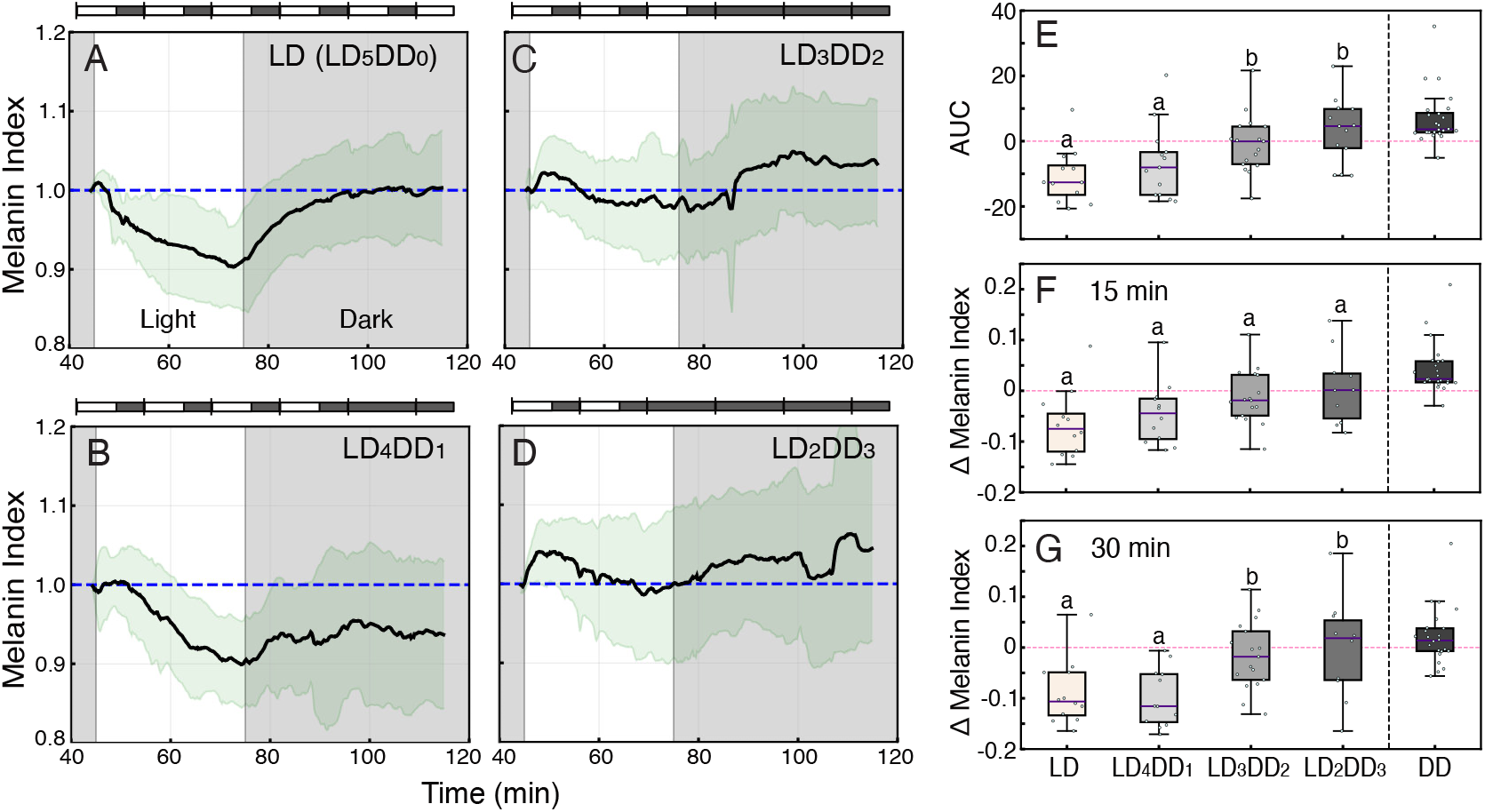
Light-induced body color changes of 5-dpf larvae reared in LD cycles followed by dark rearing. (A-D) Time courses of melanin index in 5-dpf recordings of larvae subjected to LD-rearing (A, n = 12), to four LD cycles followed by dark rearing, named LD_4_DD_1_ (B, n = 12), to three LD cycles followed by dark rearing, named LD_3_DD_2_ (C, n = 17) , and to two LD cycles followed by dark rearing, named LD_2_DD_3_ (D, n = 10). Mean values of melanin index (± SD, green shadow) are indicated. (E) Comparison of AUC among the indicated conditions. (F, G) Comparison of melanin index changes in 15 minutes (F) or 30 minutes (G) starting from light onset among the indicated conditions. DD data in panels E-G are reproduced from those in Figure 1 and excluded from the statistical comparisons. The alphabets indicated in panels E-G refer to statistical comparison using Tukey-HSD test (p < 0.05).

To investigate whether light-dependent conversion occurs following just a single exposure to light, we exposed DD-reared larvae to light just for 14 hours, from 9:00 to 23:00, on 2, 3, or 4 dpf and measured their light-induced body color change on 5 dpf (Figure 3). The larvae exposed to 14-h light on 2 dpf (2-L; Figure 3A) showed a transient darkening in response to light on 5 dpf, just like the DD-reared ones (Figure 1C). In contrast, both the larvae exposed to 14-h light on 3 dpf (3-L; Figure 3B) or 4 dpf (4-L; Figure 3C) exhibited light-induced whitening on 5 dpf, like the LD-reared ones (Figure 1B). This difference among the three conditions were supported by calculations of AUC (Figure 3D) and melanin index change during the light exposure (Figure 3E, F), which all showed statistical significant difference from 2-L to 3-L and 4-L. Note that 3-L and 4-L were not statistically different from each other in these calculations but the calculated values for 4-L looked closer to those for LD (Figure 3E-F). These results indicated that a single 14-h light exposure on 3 or 4 dpf is sufficient for conversion of body color change pattern to whitening on 5 dpf, and also suggested that light exposure on 4 dpf might be more potent for the conversion.

**Figure 3.**
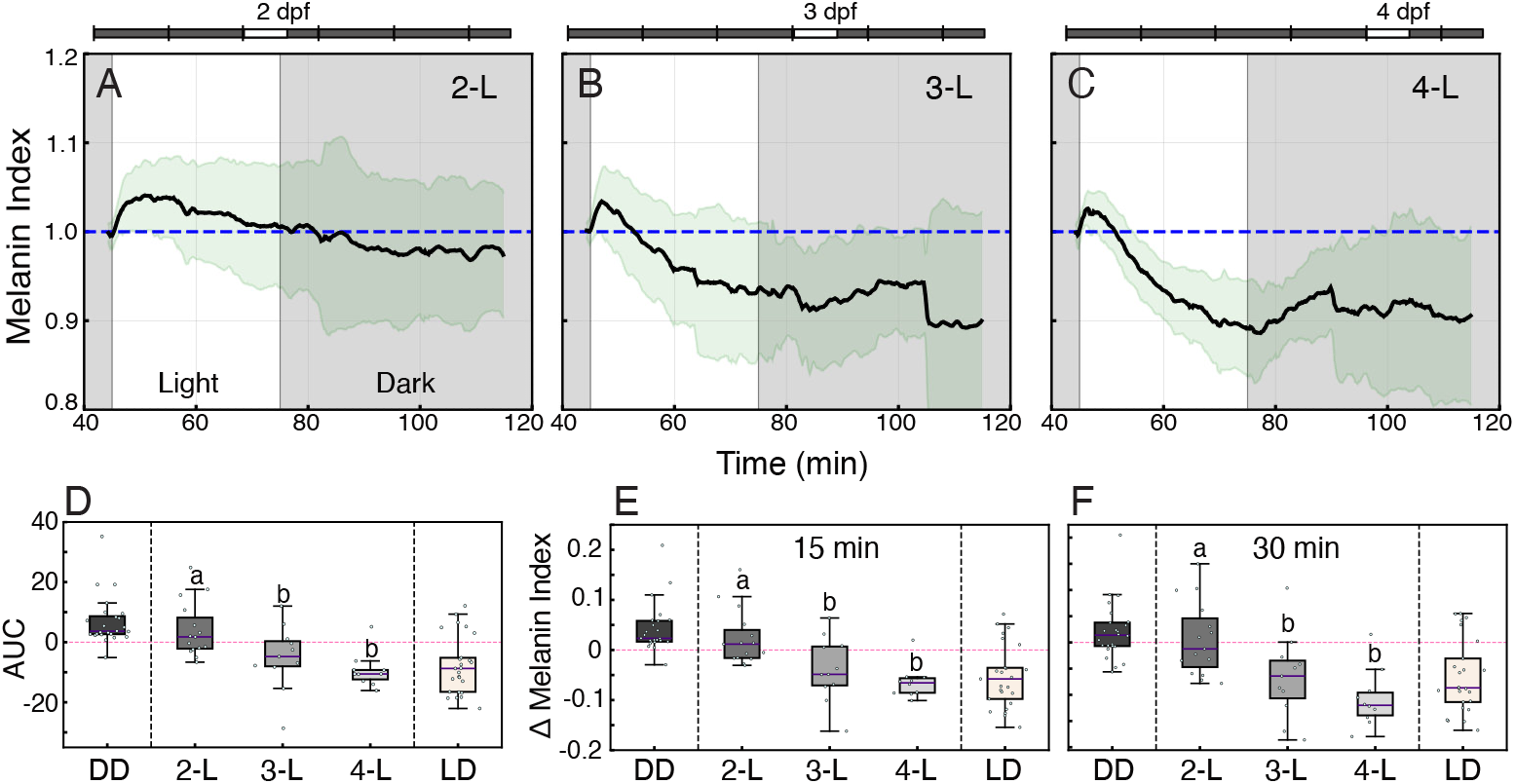
Light-induced body color changes of 5-dpf larvae reared in the dark with a single 14-h light exposure on 2 dpf (2-L), 3 dpf (3-L) or 4 dpf (4-L). (A-C) Time courses of melanin index in 5-dpf recordings of larvae subjected to 2L condition (A, n = 15), to 3L condition (B, n = 11), and to 4L condition (C, n = 11). Mean values of melanin index (± SD, green shadow) are indicated. (D) Comparison of AUC among the indicated conditions. (E, F) Comparison of melanin index changes in 15 minutes (E) or 30 minutes (F) starting from light onset among the indicated conditions. LD and DD data in panels D-F are reproduced from those in Figure 1 and excluded from the statistical comparisons. The alphabets indicated in panels D-F refer to statistical comparison using Tukey-HSD test (p < 0.05).

To further narrow down the time window inducing the conversion, we next exposed DD-reared larvae to a shorter (5 h) light illumination in the morning (Zeitgeber time [ZT] 0-5), afternoon (ZT4-9), or evening (ZT9-14) of 4 dpf (Figure 4A; ZT0 = 9 a.m.; see Materials and Methods). In all three time windows, the mean melanin index decreased during the light phase on 5 dpf, as observed in the 4-L condition (Figure 4C, D, F, H). In the quantitative comparisons, however, most of the 5-h conditions were statistically indistinguishable from both DD and 4-L, and only 4L-a (1x) in AUC (Figure 4J) and 4L-a (1x and 3x) in the 15-min melanin index change (Figure 4K) differed significantly from DD. Being concerned for accumulation of light intensity along the exposure time, we also examined the 3-fold intensity (3x) of 5-h light for each time window and obtained similar results to the 1x conditions (Figure 4E, G, I). Thus, 5-h light exposure on 4 dpf tended to induce whitening on 5 dpf irrespective of the time of day, but the present data do not allow us to conclude that 5 h of light is sufficient, or to identify a specific time window.

**Figure 4.**
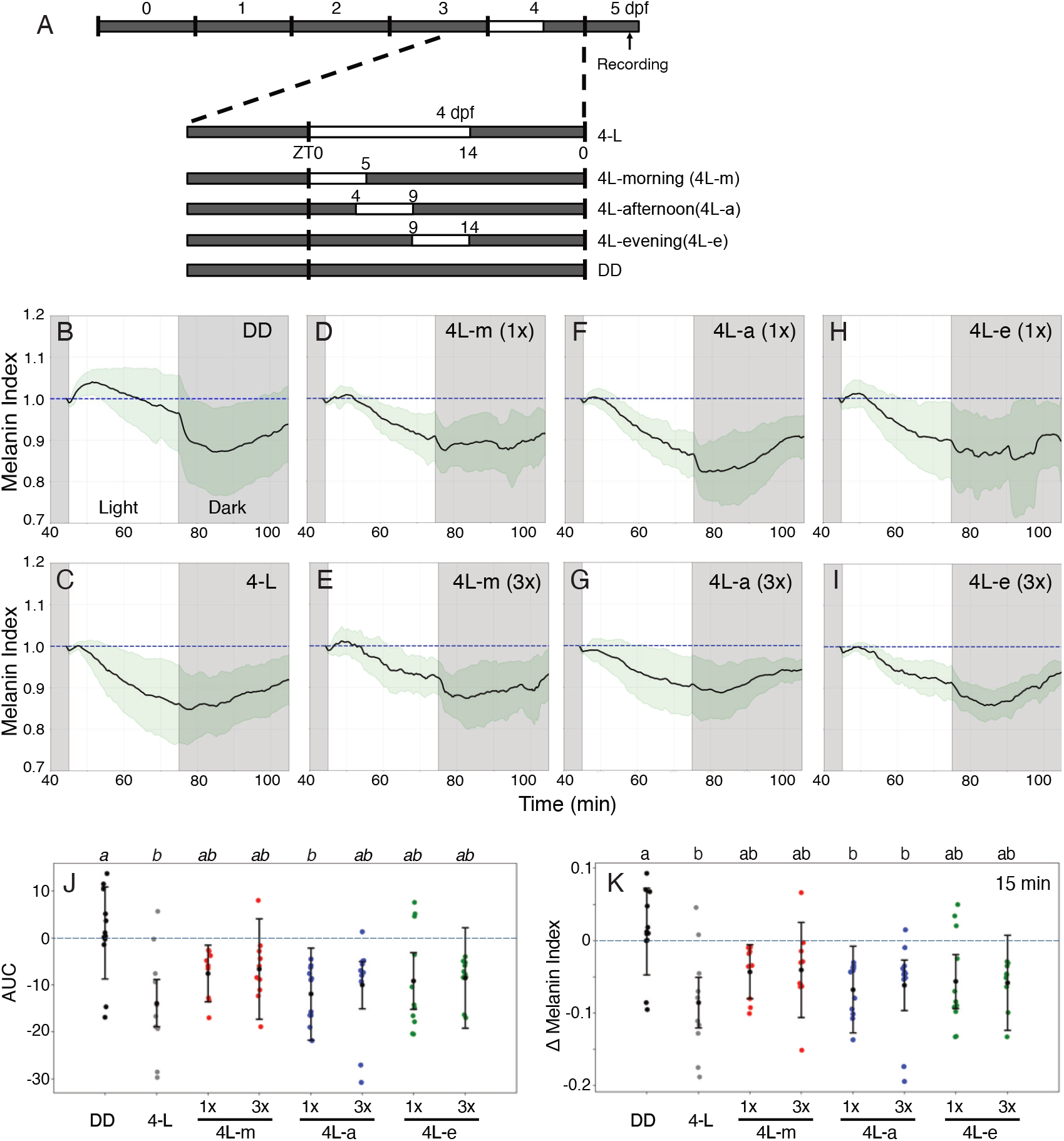
Light-induced body color changes of 5-dpf larvae reared in the dark with a single 5-h light exposure on 4 dpf. (A) The rearing strategy. The larvae were reared in the dark until 4 dpf and exposed to 5-h light in ZT0-5 (4L-morning), in ZT4-9 (4L-afternoon), or in ZT9-14 (4L-evening) and then kept in the dark again (ZT0 = lights-on at 9 a.m.). The larvae reared in the dark (DD) or exposed to a single 14-h light on 4 dpf (4L) were used for experimental comparison. (B-I) Time courses of melanin index in 5-dpf recordings of larvae subjected to DD (B, n = 11), 4L (C, n = 10), 4L-morning (D and E, n = 10 each), 4L-afternoon (F and G, n = 11 each), and 4L-evening (H and I; n = 11 and 10, respectively). Mean values of melanin index (± SD, green shadow) are indicated. In panels E, G and I, the larvae were exposed to 3-fold higher intensity of light. (J) Comparison of AUC among the indicated conditions. (K) Comparison of melanin index changes in 15 minutes starting from light onset among the indicated conditions. The alphabets indicated in the panels J and K refer to statistical comparison using Tukey-HSD test (p < 0.05).

### Retinal photoreceptor cells looked normal in dark-reared larvae

The timing of light exposure sufficient for the conversion, 3 or 4 dpf, overlaps with the developmental course of retina: Eye development begins as early as 28 hpf with the differentiation of retinal ganglion cells (Larison & Bremiller, 1990), and the typical laminar structure of retina begins to be detected at around 50 hpf and finally develops across retina by 74 hpf (Schmitt and Dowling, 1999). At 72 hpf, however, the visual function has not been completely established (Easter and Nicola, 1996), which implies that the local circuit in retina or innervation to the central nervous system is still being established. We thus examined whether light deprivation in rearing affect retinal morphology by observing the retinal section (Figure 5). The retina of DD-reared larvae had a laminar structure occupying the whole retina containing all discernible layers including ganglion cell layer, inner nuclear layer, and outer nuclear layer as observed in LD-reared larvae (Figure 5A, B). We also examined rod and cone photoreceptors in the retinal sections of 5-dpf transgenic larvae*, Tg(rho:egfp)* and *Tg(gnat2:egfp)* (Figure 5C-F): The photoreceptor cells in the retina showed no detectable difference between DD- and LD-reared larvae. The retina of dark-reared larvae looked morphologically similar to those of LD-reared larvae and exhibited no observable indication detected in the earlier-stage immature retina.

**Figure 5.**
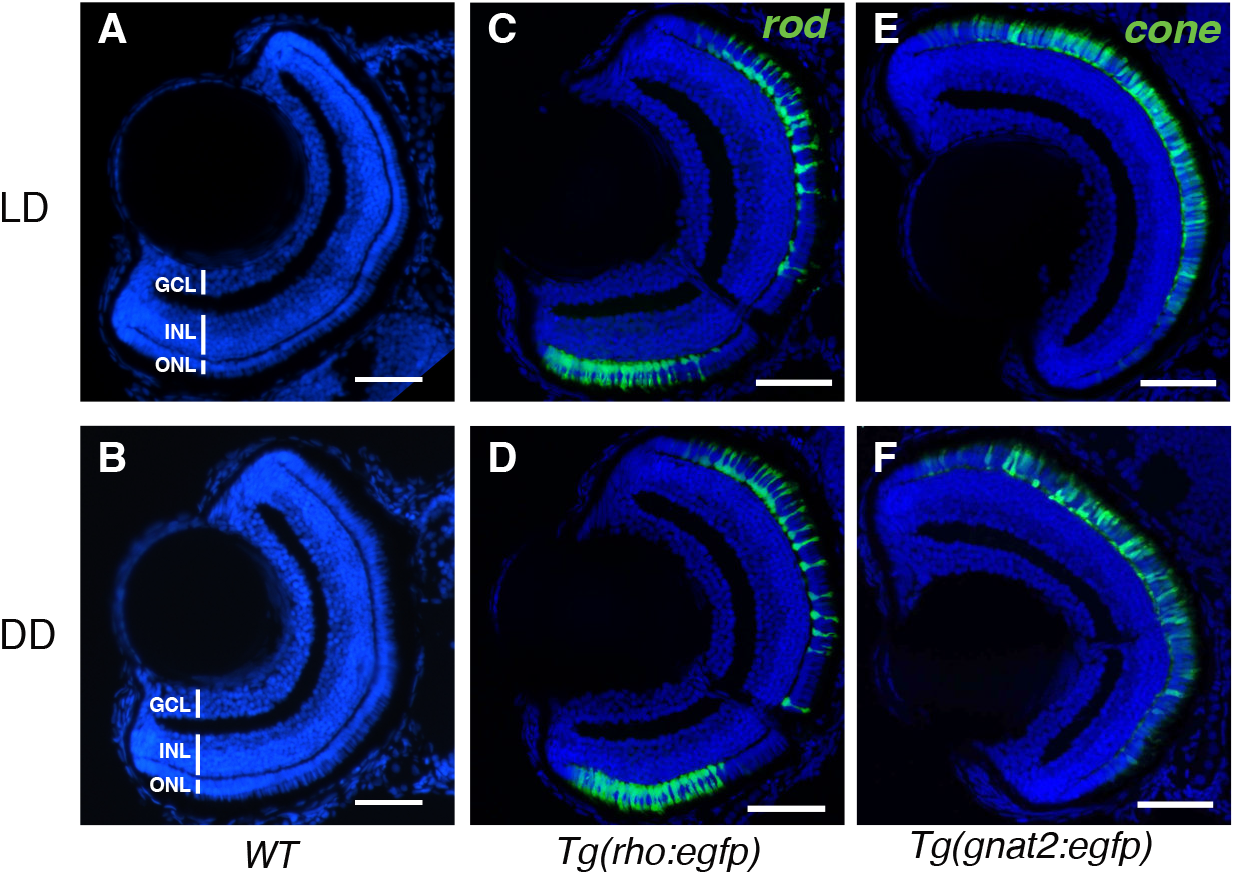
Retinal morphology and rod and cone photoreceptor cells in LD- and DD-reared larvae. (A, B) DAPI-stained retinal sections of 5-dpf LD-reared (A) and DD-reared (B) wild-type larvae. (C, D) EGFP fluorescence (green) of rod photoreceptor cells in the DAPI-stained retinal sections of 5-dpf LD-reared (C) or DD-reared (D) *Tg(rho:egfp)* larva. (E, F) EGFP fluorescence (green) of cone photoreceptor cells in the DAPI-stained retinal sections of 5-dpf LD-reared (E) or DD-reared (F) *Tg(gnat2:egfp)* larva. Scale bar = 50 μm. GCL, ganglion cell layer; INL, inner nuclear layer; ONL, outer nuclear layer.

### Pineal-specific photoreceptor cell ablation in Tg(*exorh:mCherry-2A-ntr2.0*) transgenic larvae

Although the eyes in larval zebrafish are required for light-induced whitening on 5 dpf (Shiraki et al., 2010), a previous report has suggested a role of the pineal gland, another photosensitive organ, in this light response of larval zebrafish: a mutant which do not develop pineal gland (Masai et al., 1997) fails to whiten themselves in response to light despite having morphologically normal eyes (Zhang et al., 2010). To examine a possible role of the pineal gland in light-induced conversion of body color change pattern, we next conducted specific ablation of pineal photoreceptor cells.

To this end, we applied a cell ablation technique by using nitroreductase (NTR) and its prodrug metronidazole (Mtz). We first established a transgenic zebrafish which specifically expressed a modified version of NTR2.0 (Sharrock et al., 2022) as well as mCherry in a majority of pineal photoreceptor cells under the control of ∼1kb *exorh* promoter (Asaoka et al., 2002) (Figure 6A). The resulting zebrafish line was named Tg(*exorh:mCherry-2A-NTR2*) and used for fluorescence examination and cell ablation experiments (Figure 6B). The Tg(*exorh:mCherry-2A-NTR2*) larvae showed clear red signals in the larval pineal gland on 4 dpf (Figure 6C, C’, C"; arrowheads), which successfully disappeared by treatment with 0.8 mM Mtz (Figure 6D, D’, D") to a comparable level of wild-type larvae in a similar condition (Figure 6E, E’, E"). Note that the larval treatment with 0.2–1.2 mM of Mtz for 48 h did not affect body length, indicating that this range of Mtz concentration is feasible for NTR-dependent ablation experiment. Then, the larvae that were dark-reared, pineal-ablated and exposed to 14-h light on 4 dpf (Figure 6B) were subjected to recording of the body color change pattern on 5 dpf (Figure 6F). The 4-L conditioned, pineal ablated larvae showed light-induced whitening on 5 dpf (Figure 6F) in a similar manner to the 4-L conditioned wild-type larvae (Figure 6G), whereas DD-reared wild-type control exhibited an opposite pattern (Figure 6H). In the quantitative analyses, the Mtz-treated Tg group was not significantly different from the Mtz-treated wild-type 4-L group, and differed significantly from DD only in the 15-min melanin index change (Figure 6J); in AUC and the 30-min change it was not statistically separable from either group (Figure 6I, K), probably reflecting the large variation and small sample size. These results indicate that the conversion occurred even after ablation of most *exorh*-positive photoreceptor cells, although a partial contribution of these cells cannot be excluded.

**Figure 6.**
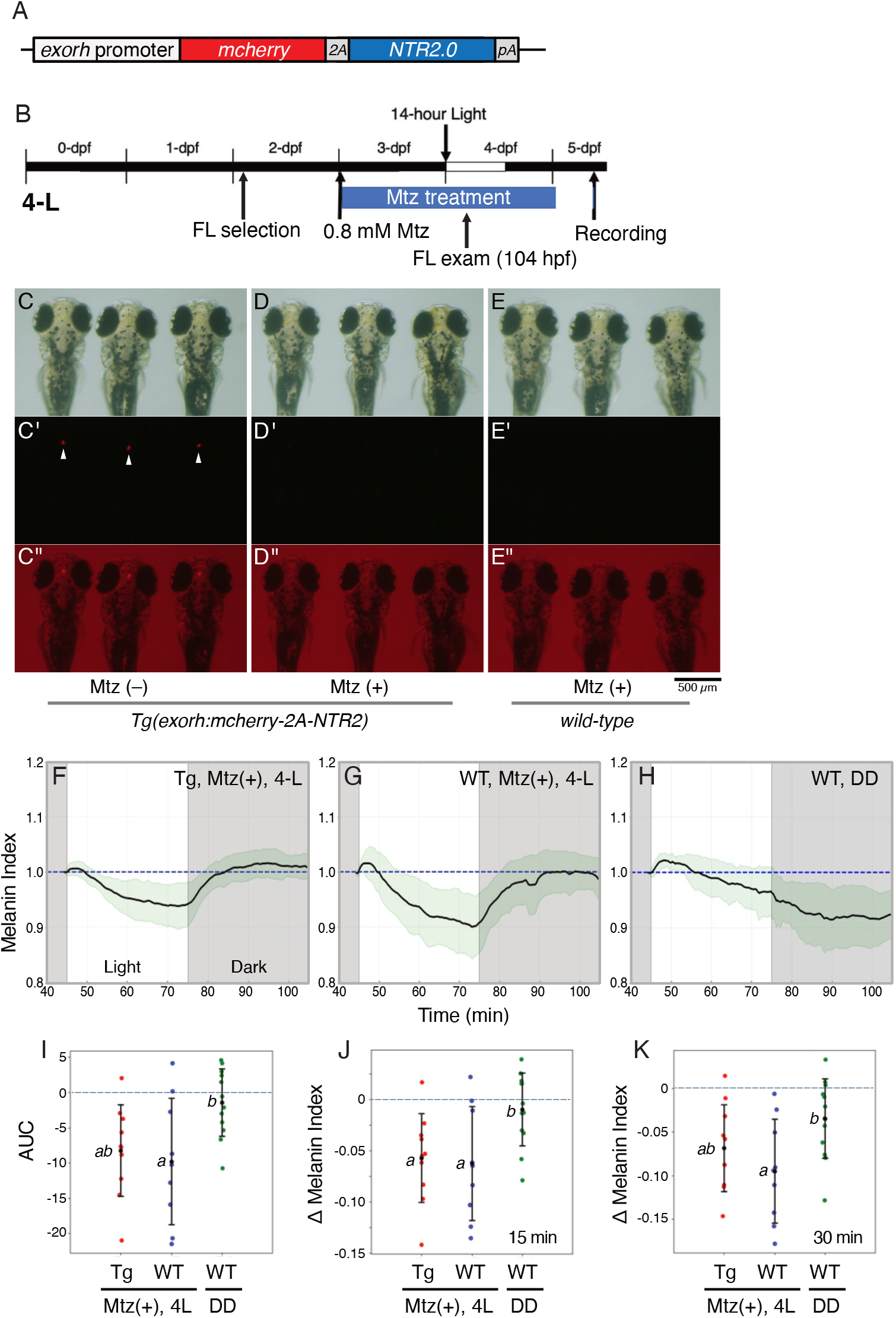
Pineal photoreceptor cell ablation and its effect on light-dependent transition of body color change pattern. (A) Schematic drawing of the transgene element used to establish transgenic line expressing NTR2.0 under control of *exorh* promoter. (B) The schedule of rearing and Mtz treatment of the *Tg*(*exorh:mCherry-2A-NTR2.0*) larvae. (C-E) Bright-field images on 4-dpf for *Tg*(*exorh:mCherry-2A-NTR2.0*) larvae (C, D) or wild-type larvae (E) without (C) or with Mtz treatment (D, E). (C’-E’) Fluorescent images of mCherry labeling. Arrowheads point to mCherry signals in the *exorh*-positive pineal photoreceptor cells (C’). (C"-E") Fluorescent signals of C’-E’ superimposed on the bright-field view. (F-H) Time courses of melanin index in 5-dpf recordings of *Tg*(*exorh:mCherry-2A-NTR2.0*) (F, n = 10) or wild-type larvae (G, n = 11) subjected to Mtz-treatment followed by 14-h light exposure on 4 dpf, or wild-type larvae reared in DD without Mtz-treatment (H, n = 10). Mean values of melanin index (± SD, green shadow) are indicated. (I) Comparison of AUC among the indicated conditions. (J, K) Comparison of melanin index changes in 15 minutes (J) or 30 minutes (K) starting from light onset among the indicated conditions. The alphabets indicated in panels I-K refer to statistical comparison using Tukey-HSD test (p < 0.05).

### Dark-reared 5-dpf larvae had a decrease in *pmch* and *pmchl* expression

In teleost, body colors could be regulated by two melanin-concentrating hormone (MCH, *pmch* and *pmchl* in zebrafish) and α-melanocyte-stimulating hormone (α-MSH, *pomca* in zebrafish). Both *pmch* and *pmchl* are expressed in separate neurons of the hypothalamus in zebrafish, and the expression is detectable as early as on 2 dpf (Berman et al., 2009). Projections from the MCH neurons are in proximity to the bloodstream, indicating that they act as not only neuropeptides but also hormones in zebrafish (Madelaine et al., 2020). Moreover, dark-adapted transgenic zebrafish, in which expression of mammalian MCH orthologue is induced by heat shock, whiten themselves under heat shock (Madelaine et al., 2020), suggesting their functions *in viv*o. Two MCH receptors, *mchr1a* and *mchr1b*, are expressed in many regions in the brain while *mchr2* is expressed in skin cells and shown to induce the pigment aggregation *in vivo* (Madelaine et al., 2020). On the other hand, *pomca* expression is localized in pituitary gland and detectable as early as 23 hpf (Hansen et al., 2003). These accumulated evidence, together with the present results on light-dependent conversion of body color change pattern in larval zebrafish, raised a question whether and how the light-rearing conditions such as DD and 4-L affect expression levels of these hormonal genes.

To answer this question, we extracted RNA from the larvae reared under DD, 2-L, 3-L, 4-L, and LD conditions and subjected them to RT-qPCR analyses (Figure 7). As a result, the DD-reared larvae had greatly reduced transcription levels of *pmch* and *pmchl* compared to LD-reared larvae (Figure 7A, B). Therefore, LD-reared larvae may have a higher capacity for MCH production than dark-reared larvae, although whether this difference accounts for the light-induced body color change remains to be determined. In contrast, *pomca* expression tended to be higher in DD-reared larvae than in LD-reared larvae (approximately 2.4-fold, *p* = 0.065), and was significantly higher in 2-L and 4-L larvae than in LD-reared larvae (Figure 7C). On the other hand, the expression levels of the three genes in the DD-reared were not significantly different from those in the larvae reared under 3-L and 4-L condition (Figure 7A-C), which were shown above to induce the conversion of body color change (Figure 3). Although the mean expression levels of *pmch* and *pmchl* in 4-L larvae were numerically higher than those in DD larvae, the difference was not significant (Figure 7A, B). Thus, *pmch*/*pmchl* expression did not correspond to the presence or absence of the light-induced whitening.

**Figure 7.**
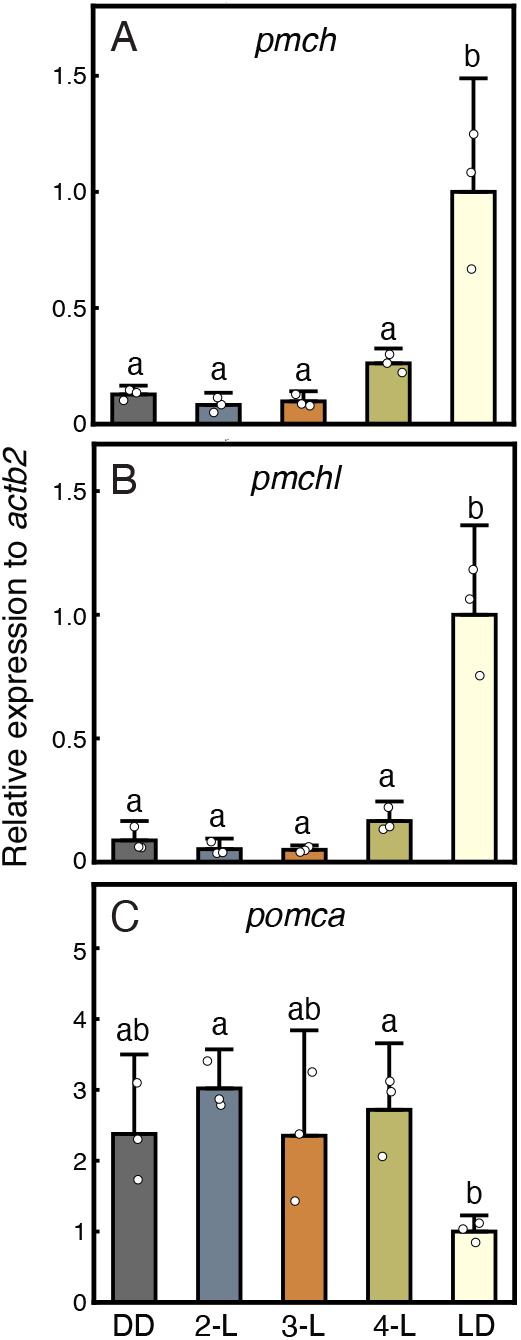
Gene expression differences in signaling molecules for body color change. The transcription levels of *pmch* (A), *pmchl* (B) or *pomca* (C) were normalized to *actb2* expression level and compared among the larvae reared in the indicated light conditions (n = 3 biological replicates per condition; each replicate is a pool of 12 larvae). The larvae were sampled at ZT6 (3 p.m.) on 5 dpf. Bars and error bars indicate mean ± SD, and open circles indicate individual replicates. The alphabets indicated in panels A-C refer to statistical comparison using Tukey-HSD test (p < 0.05).

## Discussion

### Dark-reared 5-dpf larvae show a "immature" pattern of body color change, which is rescued by a single light exposure on 3 or 4 dpf

LD-reared larvae, as shown in Figure 1, whitened themselves in response to light on 5 dpf. In contrast, dark-reared larvae darkened themselves in response to light, resulting in a darker body color during the light phase (Figure 1C). This phenomenon reminds of the light-induced darkening of body color change in 2-dpf larvae (Shiraki et al., 2010), but it is not completely the same. LD-reared 2-dpf larvae darken themselves and keep darkened color at least for 30 min (Shiraki et al., 2010), while dark-reared 5-dpf larvae darken themselves only in the first 10 minutes and then gradually whitening (Figure 1C). One possible explanation is that the light-induced darkening observed in 2-dpf larvae may reflect lack in the critical neural circuit controlling body color change, whereas such circuit may be present but immature (or incomplete) in the 5-dpf dark-reared larvae, unveiling the important role of light in shaping the neural circuit. In fact, it has been shown that light or visual experience helps shape the visual function in zebrafish (Avitan et al., 2017; Reynolds et al., 2026). Although previous researchers did not find any difference in the innervation between the eye and the optic tectum of dark-reared larvae (Niell and Smith, 2005), the advanced technique and statistical analysis detected modest changes in the pattern of spontaneous retinal activity, in network dynamics of optic tectum and in feeding behavior of larval zebrafish dark reared (or vision-deprived) from 5 dpf to 8 dpf (Avitan et al., 2017). Similarly, in dark-reared larvae, retinal PSD-95 protein density was lower (Xie et al., 2019). These studies highlight the role of light in shaping neural network and in modifying visual ability. Intriguingly, our current study demonstrated that a single light exposure on 3 dpf or 4 dpf also converted the body color change pattern on 5 dpf in dark-reared larvae (Figure 3), suggesting the presence of "critical period" in the maturation of body color change ability. The critical period refers to a time interval when some external stimuli are absolutely required for developmental changes in certain neural circuits and normal abilities and phenotypes (Hooks and Chen, 2007). For example, visual experience at early stage is required for the formation of ocular dominance in kittens (Hubel and Wiesel, 1970). Future experiments giving a single light exposure to dark-reared larvae at the later stages would help us examine the presence of time window similar to the critical period.

One possible scenario to be considered on the reversed or "immature" pattern of body color change in dark-reared larvae (Figure 1) might be that it stems from or is secondary to the immature circadian rhythm. Some research pointed out that the circadian rhythm in cellular level, specifically the oscillation of *per1* transcription, arises intrinsically and is detectable on the first day after birth of zebrafish embryos (Dekens and Whitmore, 2008). However, in tissue or embryonic level, there is no rhythmicity in mRNA of *per1* or *per2* in dark-reared larval zebrafish (Dekens and Whitmore, 2008), which reflects the desynchronization of cell population. Additionally, melatonin rhythm was found to be absent in dark-reared larvae (Kazimi and Cahill, 1999). Since melatonin plays multiple roles *in vivo*, dark rearing might bring about physiological dysfunctions (de Borsetti et al., 2011) . In behavioral level, most of larvae reared in darkness lose the locomotor rhythmicity (Hurd and Cahill, 2002). Therefore, the entrainment by light/dark cycles is indispensable for the establishment of circadian rhythm. Much importantly, light itself is enough for the entrainment as a 1-hour light exposure within 24 hpf initiates the oscillation of transcripts of clock genes in individual level and the synchronization of mRNA expression of clock proteins in tissue is promoted after light exposure (Dekens and Whitmore, 2008). Light before 14 hpf is incapable to initiate the melatonin rhythm in dark-reared larvae (Kazimi and Cahill, 1999) while light from 14 hpf to 24 hpf is capable (Ziv and Gothilf, 2006). Yet, the locomotor rhythmicity does not exist in larvae receiving light from 0 hpf to 14 hpf (Hurd and Cahill, 2002). Hence, in zebrafish hardwires for circadian rhythm are important, but light, as an external environment cue at an appropriate timing, is also essential. In our experiment, the anomalous darkening observed in dark-reared larvae may be secondary to the unsynchronized circadian rhythm, but the following experiments gave some indirect evidence to disproving it. In Figure 2, we initially gave all larvae with light/dark cycles, which are expected to synchronize the circadian rhythm as the LD group, but they did not show clear whitening on 5 dpf, although the data was of huge variation. In Figure 3, a period of light was given on 2, 3 or 4 dpf, and as shown in previous study, this duration of light would synchronize cellular clock and establish the circadian rhythm. From this viewpoint, despite of the larvae in Figure 3 probably having the normal circadian rhythm, one of them (2-L) failed to whiten themselves in response to light, suggesting the deficit in light-induced body color change was not secondary to disrupted circadian rhythm in dark-reared larvae.

### Possible photoreceptive mechanisms regulating the conversion of body color change

Our current study demonstrated that a single light exposure on 3 or 4 dpf is sufficient for conversion of body color change ability (Figure 3), raising a question which photoreceptive cells are responsible for this light-dependent conversion. Larval zebrafish have two major photosensitive tissues, retina and pineal gland, which start and finish development in different periods. The pineal gland is formed by 20 hpf (hours-post-fertilization) (Wilson and Easter, 1991; Masai et al., 1997) and the photoreceptor markers become detectable in pineal by 24 hpf (Heisenberg et al., 1996). At the same time, melatonin rhythm is detectable (Kazimi and Cahill, 1999). On the other hand, eye development begins at 28 hpf, with the appearance of retinal ganglion cells (Larison and Bremiller, 1990). The development course is quite slow compared to pineal: the photoreceptors are detectable by 48 hpf (Raymond et al., 1995), the connection to optic tectum is fully built on 72 hpf (Burrill and Easter, 1994) and the vision ability is matured at about 80 hpf (Easter and Nicola, 1996). Therefore, developmental time windows for both the pineal gland and the retina may fit the initial timing of light (3 dpf) when it can induce conversion of body color change ability. Although a previous study (Zhang et al., 2010) suggested a potential role of the pineal gland, pineal-specific ablation did not abolish the light-induced conversion by a 14-h light on 4 dpf (Figure 6), arguing against a major contribution of pineal *exorh*-expressing photoreceptive cells. A partial contribution of these cells, however, cannot be excluded, because the Tg group was not statistically separable from DD in AUC and the 30-min melanin index change (Figure 6I, K). It should be noted that many photoreceptor cells express *exorh* but others express red cone opsin, parietopsin or parapinopsin1 (Mano et al., 1999; Wada et al., 2018; Cau et al., 2019; Sapède et al., 2020) and they may have a role in this light-dependent conversion.

As for the retina, we briefly examined the larval retinal sections for the layered structure as well as rod and cone photoreceptor cells, but did not observe any obvious abnormality in the dark-reared retina (Figure 5). Previous study reported that larvae lacking visual photoreceptors (cones and rods) are still capable of body color change to light (Muto et al., 2005), while the larvae lacking retinal projection to brain are unable (Kay et al., 2001). These evidence suggest a potential role of non-rod non-cone photosensitive cells in the retina. Zebrafish genome has five melanopsin genes (*opn4m1, opn4m2, opn4m3, opn4x1* and *opn4x2*), some of which contribute to a role in increasing locomotor activity in the wake state (Dekens et al., 2022) and in the circadian locomotor rhythm (Chaigne et al., 2024). Their transcripts are all detectable in 3 to 5 dpf larval retinas and the transcription of opn4x1 and opn4m2 was found to decrease under a longer photoperiod (18-h light/6-h dark cycle) (Matos-Cruz et al., 2011). It is worthwhile exploring possible contribution of the melanopsin-expressing neurons in retina in terms of light-dependent conversion of body color change pattern. Because retina has a quite sophisticated neural circuit, possible abnormality in interneurons of dark-reared retina should be taken into account. For example, dopaminergic neurons adopt the role of elevation of visual sensitivity in zebrafish (Li and Dowling, 2000). The transcription of tyrosine hydroxylase (*th*) could be detected as early as 96 hpf in retina (Holzschuh et al., 2001). Interestingly, larval zebrafish reared in 100 μM fluphenazine (an antagonist for dopamine D2 receptor) from 2 hpf exhibit a smaller eye size and no light-induced whitening in 15-min light exposure (Syambani Ulhaq, 2020), being reminiscent of our results. It is intriguing to investigate the number of dopaminergic neurons and dopamine D2 receptor expression in dark-reared larvae. What is equally important is the innervation from retina. Larval retinal projection reaches optic tectum and many other regions. Intriguingly, it was reported that retinal ganglion cells sent axons to the lateral hypothalamus which resided the *pmch* and *pmchl* neurons and that some of those retinal ganglion cells expressed *opn4x1*, suggestive of a possible and direct pathway from light sensing to hormone control in body color change (Kölsch et al., 2021; Slanchev et al., 2025). Thus, it is interesting to examine the innervation of retina in dark-reared larvae.

### *Pmch* and *pmchl* transcription were in lower level in dark-reared larvae

The current study found out that *pmch* and *pmchl* transcription was in lower levels in dark-reared larvae than in LD-reared ones (Figure 7), which is in line with the experiments using larvae reared in white or dark background (Berman et al., 2009; Zhang et al., 2010) although the reported ratio (∼30 %) was not as low as those in this study (∼10%, DD vs LD ; Figure 7A,B). Because the experimental paradigms differ, these values should not be compared directly. Among the groups that received a single light exposure, the mean pmch and pmchl levels in 2-L larvae were 35% and 40% of those in 4-L larvae, respectively, but these differences were not statistically significant (Figure 7A, B; Tukey-HSD). In addition, 3-L and 4-L larvae, which showed the conversion to whitening, were not significantly different from DD larvae, whereas 4-L larvae remained below the LD level (Figure 7A, B). Given the small sample size (n = 3), we cannot exclude a modest effect of light exposure on *pmch/pmchl* expression, but our data do not show that the level of these transcripts corresponds to the presence or absence of the conversion. Could the diminished *pmchl* and *pmch* expression account for the observed darkening in light phase in the dark-reared larvae? Endocrine pathway is always considered to be slow while the darkening in the dark-reared larvae was held for the first 10 minutes in light (Figure 1). To our knowledge, there has been no preceding study investigating the time course of MCH release and its work, except for one report showing that *in vitro* application of MCH to zebrafish melanophores causes aggregation of melanosomes (Logan et al., 2006): When the MCH is applied at 0.1 nM, it takes less than 15 minutes for melanophores to exhibit up to 50% maximal aggregation. Therefore, at present, the possibility that the endocrine pathway is involved in light-induced body color change shown here cannot be excluded. Further investigation should figure out the time course of MCH release and work by ELISA or using their reporter genes if feasible.

The larvae reared in black background (that is, the ground is black and LD cycles exist) also showed a decreased *pmch* and *pmchl* level (Zhang et al., 2010). In our experimental setting, we did not prepare the black background group. If the darkening in light on 5 dpf is attributed to the decreased *pmch* and *pmchl* level, the larvae reared on black background may be expected to exhibit the darkening or postponed whitening as the DD group above since those larvae also had a decreased level. In the future, this experiment is interesting and gives us the hint whether the decreased hormone level is the only reason for the darkening to light.

In conclusion, light in rearing is indispensable for the conversion of body color change in zebrafish. Dark-reared larvae darkened themselves on 5 dpf facing light, instead of whitening as LD-reared larvae. Light on 3 or 4 dpf could elicit the conversion whereas light before that failed. From this timing, the retinal development was predicted to be disrupted by lack of light in rearing, but there was no obvious difference in retinal morphology and rod and cone photoreceptor cells. More experiments targeting other photosensitive cells in retina should be put on the future schedule. In the downstream, we detected the diminished transcription of *pmch* and *pmchl*, the two genes encoding melanin concentrating hormones in zebrafish, which was, however, not associated with the conversion in 3-L and 4-L larvae and deserves further investigation.

## Material and Methods

### Experimental animals

Ekkwill strain was used as wild-type zebrafish. To obtain embryos, a pair of female and male zebrafish were placed in a tank before the day of mating. The mating began at 9 a.m. (counted as 0 hour-post-fertilization (hpf)) and lasted for 1-2 hours. The spawned eggs were collected and partitioned evenly into multiple dish filled with egg water (0.15% artificial sea water). For dark (DD) rearing, the fertilized eggs were placed into a light-tight box in the incubator with a setting temperature 28.5 °C. For light/dark cycle (LD) rearing, they were placed in a transparent box in the same incubator. In the incubator, the daily light period was set from 9 a.m. to 11 p.m. (14 h-light/10 h-dark cycles). In this study, Zeitgeber time (ZT) 0 was defined as 9 a.m., the time of lights-on in the incubator, which was also the time of fertilization (0 hpf); thus, ZT0–14 corresponds to the light period and ZT14–24 to the dark period of the LD cycle. For DD-reared larvae and for larvae kept in the dark during the dark-rearing periods, ZT was defined in the same manner, assuming a 24-h cycle starting from 9 a.m. (0 hpf), although these larvae did not experience any light/dark transitions. For those that were exposed to light or darkness at certain period, they were transferred to the transparent or light-tight box immediately before 9 a.m. when a new light/dark cycle was about to begin. The size of those boxes was 22 cm (L) x 16 cm (W) x 6 cm (H). The light source of incubator was a white LED with power of about 0.36 mW/cm^2^ on the box as the standard condition (1x).

### Measurement of body color changes

The devices for measurement of body color changes of larval zebrafish will be published in detail elsewhere. In brief, ∼1 mm-thick gel of 1% agarose (Nakalai Tesque) in egg water made in a 35-mm Φ dish was used to produce a platform with two or three troughs [∼5 mm (L) x ∼0.56 mm (W) x ∼0.56 mm (H)] in the center. The troughs were filled with 2% methylcellulose. The larvae reared in the same condition were placed in the troughs, covered by 1.5% agarose XP (NIPPON GENE) which was melt and kept at 42 °C in advance, and they were kept for ∼15 min at room temperature (∼25°C) so that the covering gel was solidified. Finally, ∼1 mL of egg water were placed above. Note that the whole gel was transparent allowing light to reach the larvae in measurement, and that the larvae (except for LD-reared ones) were kept in the dark or under deep red light (730-nm LED) during the preparation. The imaging devices including a dissection microscope (Leica S8APO) equipped with a CCD camera (Leica, DFC3000G) were placed in a black light-tight cabinet to exclude ambient light. Infrared light (∼810 nm) was used to illuminate the larvae from underneath and detected by the camera through an IR-selected filter (IR-80, Fujifilm), so as to make them visible even in the dark. The temperature in the cabinet was maintained at 27.5-29.5 °C during the measurement. The dish holding the larvae were set on the microscope stage inside the cabinet and subjected to the measurement: In a typical experiment, the larvae were subjected to a 45-min dark period, a 30-min light period and then a 45-min dark period. In the light period, the larvae was illuminated from underneath with white light from a halogen lamp (PCS-NHF 150, Nippon PI) with power of 1.5 mW/cm^2^ at the specimen. During the measurement, an image of the larvae was captured by the camera every 10 seconds to give a series of pictures in gray scale. The larvae that moved around in the trough during the measurement and/or that were found dead at the end of measurement were excluded from the following analyses. The measurements were conducted during the light phase of fifth day post-fertilization (typically between 11 a.m. and 8 p.m. [ZT2-11]).

After compiling these time-lapse images and saving them as a QuickTime movie, they were converted to a series of TIFF images using ImageJ. From these image data, the body color for each frame was quantified as follows. First, all frames were binarized using a fixed threshold so that at this value the area occupied by melanin pigment within melanophores may be appropriately displayed as black pixels. The same value was used for all measurements and all frames. After calculating the total number of black pixels throughout the body in each frame, the value following the initial dark adaptation in each experiment (45 minutes from the beginning of the recording) was set to 1.0, to which the total number of black pixels in each frame was normalized.

The normalized value was defined as the Melanin Index (MI) so as to represent the blackness of the body color.

To quantify the light-induced body color change, MI time courses were smoothed with a centered moving average over 7 frames (the frame itself and 3 frames before and after, i.e., ±30 s, corresponding to a 1-min window). After smoothing, each time course was divided by its smoothed MI value at light onset so that the baseline was 1.0, and two metrics were calculated from these values. The primary metric was the area under the curve (AUC) over the 30-min light period, defined as the area between the MI curve and the baseline (MI = 1.0). The AUC was calculated by summing (MI − 1.0) over all frames in the light period (one frame every 10 s; 181 frames), so that values above the baseline were counted as positive and those below as negative. A positive AUC therefore indicates light-induced darkening, and a negative AUC indicates light-induced whitening. As secondary metrics, we used the change in melanin index (ΔMI), defined as the difference between the MI at 15 min or 30 min after light onset and the baseline value of 1.0 (ΔMI = MI(t) − 1.0; t = 15 or 30 min).

Negative ΔMI values indicate whitening and positive values indicate darkening. AUC was designated as the primary endpoint, and ΔMI at 15 and 30 min were analyzed as supporting measures.

### Cryosection and staining

Frozen tissue sections of larval zebrafish were prepared as described previously (Kojima et al., 2008) with some modifications. In brief, larval zebrafish were anesthetized in 0.1% MS-222, and fixed in 4% paraformaldehyde (PFA) in Ca^2+^- and Mg^2+^-free Dulbecco’s PBS (D-PBS) by rotating for 2 h at room temperature. The fixed larvae were then washed three times in 5% (w/v) sucrose in D-PBS for 30 min each at room temperature, and then placed in 20% (w/v) sucrose in D-PBS at 4 °C overnight. Finally, the larvae were placed in a 1:2 mixture of Optimal Cutting Temperature Compound (O.C.T.) and 20% (w/v) sucrose solution and rotated for 60 min. The larvae were frozen in the 1:2 mixture by using liquid nitrogen so as to make a frozen cubic block containing each larva. The frozen blocks were stored at - 80 °C until cryosection. The blocks were subjected to cryosection using Leica CM3050S cryostat system to make 10-µm thick tissue slices mounted on glass slides. The slides were dried overnight at room temperature in the dark.

For nuclear staining, the cryosections on the slides prepared above were washed in D-PBS containing 0.1% Triton X-100 once and in D-PBS twice for 15 min each, and treated with 1.5 μg/ml DAPI in D-PBS for 60 min in the dark. The sections were then washed in D-PBS twice for 10 min each, coverslipped with VectaShield (Vector Laboratories) and observed with a confocal laser scanning microscope (SP8, Leica).

### RT-qPCR

Larvae at ZT6 on 5 dpf were given 45-min dark adaptation and then placed in 1.5 ml RNAlater (Sigma-Aldrich) overnight at 4 °C. After removal of RNAlater, the larvae (12 individuals each per tube) were stored at -80 °C until the following RNA extraction. Each tube containing the 12 larvae was subjected to RNA extraction (as a single sample) using RNeasy kit (Qiagen). Each pool of 12 larvae was treated as one biological replicate, and three biological replicates (n = 3) were prepared from the same clutch for each rearing condition. The purified RNA, whose concentration was measured NanoDrop One (ThermoFisher), was preserved at -80 °C. For each sample, 1 µg of RNA was subjected to cDNA synthesis and qPCR by using GoTaq 2-Step RT-qPCR System (Promega). An equal amount of cDNA from every sample was mixed together to obtain the standard cDNA sample for qPCR. PCR primers for three target genes (*pomca, pmchl* and *pmch*) and a reference gene (*actb2*) were listed in Table 1. Each qPCR reaction employed the cDNA corresponding to 0.5 ng RNA (*actb2*) or 5 ng RNA (the others). The qPCR reaction was carried out in the platform of StepOne Plus RT-PCR System (Applied Biosystem).

**Table 1.**
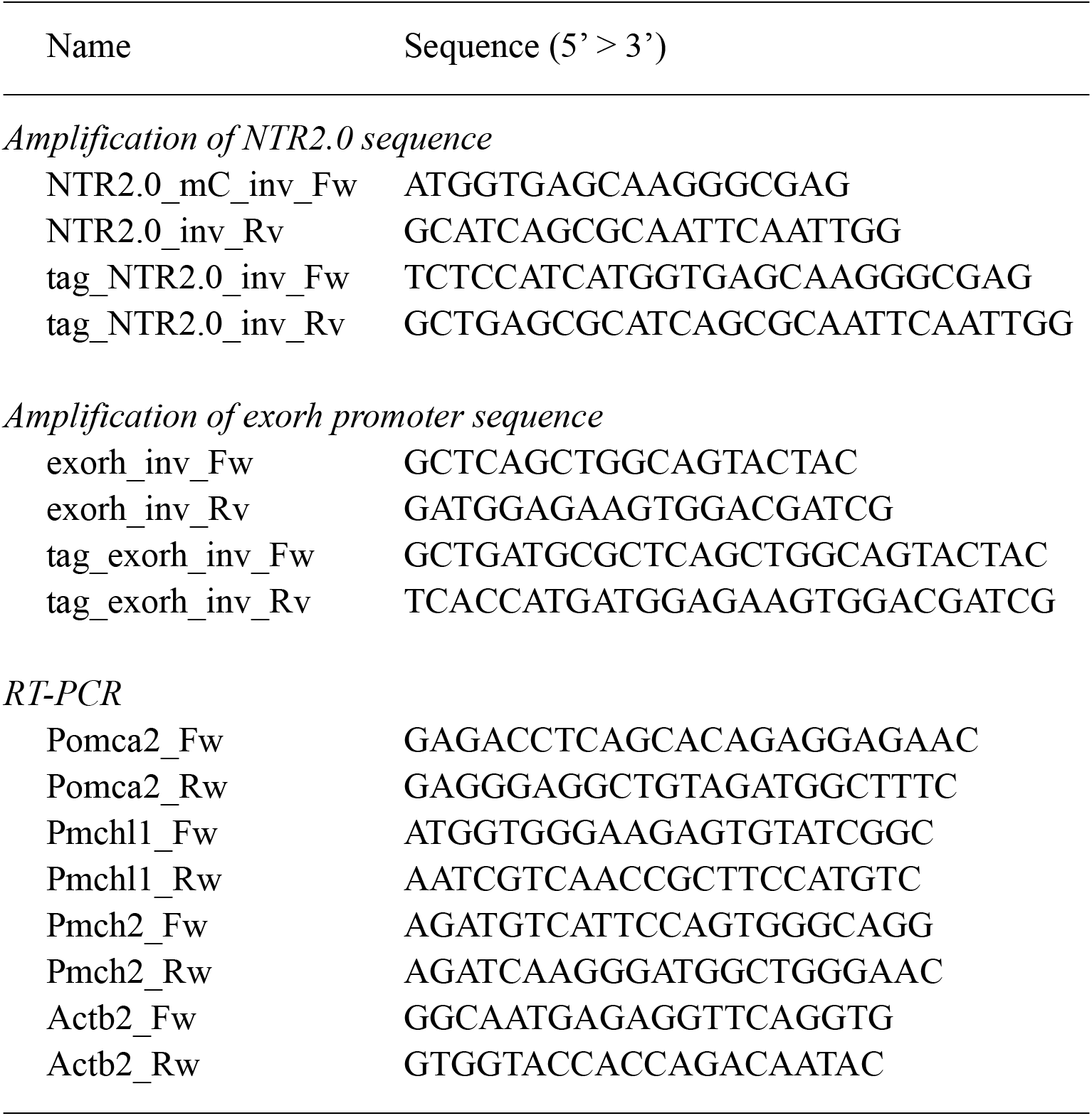
Primers used in this study.

### Establishment of transgenic zebrafish

A 1055-bp regulatory sequence of *exorh* gene (*exorh* promoter) (Asaoka et al., 2002) was used for pineal-specific expression of *NfsvB_Vv* F70A/F108Y (NTR2.0), an improved version of nitroreductase which convert the prodrug metronidazole (MTZ) into a cytotoxic product in a high efficiency (Sharrock et al., 2022). For transgene construction, the *exorh* promoter was PCR-amplified from Ex(-1055) plasmid (Asaoka et al., 2002), while a DNA fragment, containing mCherry, P2A, NTR2.0, polyA signal and Tol2 elements, was PCR-amplified from the 5xUAS:GAP-mCherry-P2A-NfsB_Vv F70A/F108Y, he:tagBFP2 plasmid (Addgene, Plasmid #158653). These DNA fragments were fused together by using In-Fusion HD Enzyme (TaKaRa), and cloned into DH-5α *E.coli*. The obtained plasmid (4 ng/μl) as well as Tol2 mRNA (25 ng/μl) were microinjected into each of one-cell stage zebrafish embryos. The injected embryos were raised to adulthood, and mated with wild-type zebrafish to obtain their offsprings, which were examined for mCherry fluorescence in the pineal gland on 5 dpf. Among them, the pineal fluorescence-positive individuals were used to establish the transgenic lines: *Tg(exorh:mCherry-P2A-NfsB_Vv F70A/F108Y,he:tagBFP2)^ja215Tg^, Tg(exorh:mCherry-P2A-NfsB_Vv F70A/F108Y,he:tagBFP2)^ja216Tg^* and *Tg(exorh:mCherry-P2A-NfsB_Vv F70A/F108Y,he:tagBFP2)^ja217Tg^*. Among these lines, the *ja216Tg* was used for the following experiment and referred to as *Tg*(*exorh:mCherry-2A-NTR2.0*) in short.

### MTZ-induced cell ablation

Hemizygous *Tg*(*exorh:mCherry-2A-NTR2.0*) was mated with wild-type zebrafish to obtain embryos. On 2 dpf, they were examined for pineal fluorescence by using a fluorescence dissection microscope (MZ-FLIII, Leica) and divided into transgene-positive and negative population. They were then transferred into 25ml egg water with MTZ for 48 hours from 72 hpf, and transferred back to 25ml egg water without MTZ for the following experiments of body color measurement on 5 dpf. For the experiment to test a dose-dependency of MTZ for pineal cell ablation, the larvae on 4 dpf were anesthetized, held in a agarose gel, and subjected to fluorescent imaging. In each image, the ROI covering a corresponding area of the pineal gland was used to calculate the grey scale value of red channel as the fluorescent intensity. This value was subtracted with the mean value of the control group (transgenic larvae raised in the absence of MTZ) to obtain the relative intensity.

### Data analysis

Data analysis and plotting were done in PyCharm and packages of matplotlib and scipy. Statistical comparisons were performed with Welch’s *t*-test between two groups, or with Tukey-HSD test among three or more groups. Image processing was done with Image J and PhotoShop. AUC was the primary endpoint; ΔMI values at 15 and 30 min were analyzed as secondary measures, and no correction was applied across these three metrics.

